# The impact of non-lineage defining mutations in the structural stability for variants of concern of SARS-CoV-2

**DOI:** 10.1101/2023.06.22.546079

**Authors:** Yasmmin Martins, Ronaldo Francisco da Silva

**Affiliations:** Data science, yPublish, Porto ALegre-RS, Brazil; Department of Pathology, Stanford University, Stanford-CA, USA

## Abstract

**Motivation:** The identification of the most important mutations, that lead to a structural and functional change in a highly transmissible virus variants, is essential to understand the impacts and the possible chances of vaccine and antibody escape. Strategies to rapidly associate mutations to functional and conformational properties are needed to rapidly analyze mutations in proteins and their impacts in antibodies and human binding proteins.

**Results:** Comparative analysis showed the main structural characteristics of the essential mutations found for each variant of concern in relation to the reference proteins. The paper presented a series of methodologies to track and associate conformational changes and the impacts promoted by the mutations.

**Supplementary information:** Supplementary data are available at *Bioinformatics* online.

## 1 Introduction

Since the emergence of the Severe acute respiratory syndrome coronavirus 2 (SARS-CoV-2) responsible for the coronavirus disease 2019 (COVID-19), a huge set of mutations have been identified in the virus genome, diversifying the original virus in multiple lineages and resulting in insertion, deletion or/and amino acid changes in the derived proteins.

Currently, five lineages originate variants of concern (VOCs), which are B.1.1.7 (alpha), B.1.351 (Beta), P.1 (Gamma), B.1.617.2 (Delta) and B.1.1.529 (Omicron), according to World Health Organization. These variants of concern are known by aggregating mutations to specific proteins that increases their fitness, changes structural properties, and consequently turns the virus more lethal and transmissible ([31]). The emerging of a new variant of Concern usually causes the vanishing of other current lineages, taking the Brazilian VOC Gamma as example, the lineages B.1.1.33 and B.1.1.28 rapidly ceased to figure in the cases [6].

Recent works have made an overview about the implications of the mutations phylogenetically determined for each VOC ([10, 31, 36]), such as response to the main vaccines, efficacy of neutralizing antibodies, review of known implications in the virus capability of transmission and binding to human receptor. These studies lack the identification and annotation of structural characteristics by *in-silico* methods, that could be used to track the behavior for future variants.

In this paper, we propose a complementary analysis to identify characteristic and essential mutations aside those that defines the VOCs. We also present a structural and functional analysis of each group of mutations according to predicted properties derived from tertiary and secondary structure,

## 2 Methods

### 2.1 Feature selection to find essential mutations for VOCs

The data used for feature selection (represented in Figure 1 A) was retrieved from GISAID containing all proteins from 603170 genomes (GISAID episet EPI_SET_20211029uv) of the lineages B.1.1.7, B.1.351, P.1, B.1.617.2 and B.1.1.529, from 2021-01-01 to 2022-01-11. The feature selection was performed among the five variants of concern (alpha, beta, gamma, delta and Omicron) and used the protein mutations. We aligned the protein sequences release with their respective SARS-CoV-2 reference proteins, and extracted the protein mutations for each sample and protein type. These mutations were represented as numerical features in a sparse binary matrix, in which 1 and 0 correspond to the presence or absence of each mutation in a certain sample, respectively. According to the data mining principle of removing the feature columns that have no variation of values in it ([27]), we removed the mutations columns that were common for the five VOCs and separated than as a group named intersection. The remaining mutation features were used to form five datasets having two classes, in the first class of these datasets there was samples of the VOC, and the rest of the samples were from the other three VOCs. The goal was the identification of the mutations that could separate one VOC from the others. We applied the Boruta method ([17]) to execute the feature selection, we applied 100 rounds containing randomly chosen four thousand samples for each class. The mutations that were classified by Boruta in rank 1 of importance in the all rounds were kept as result.

**Fig. 1.**
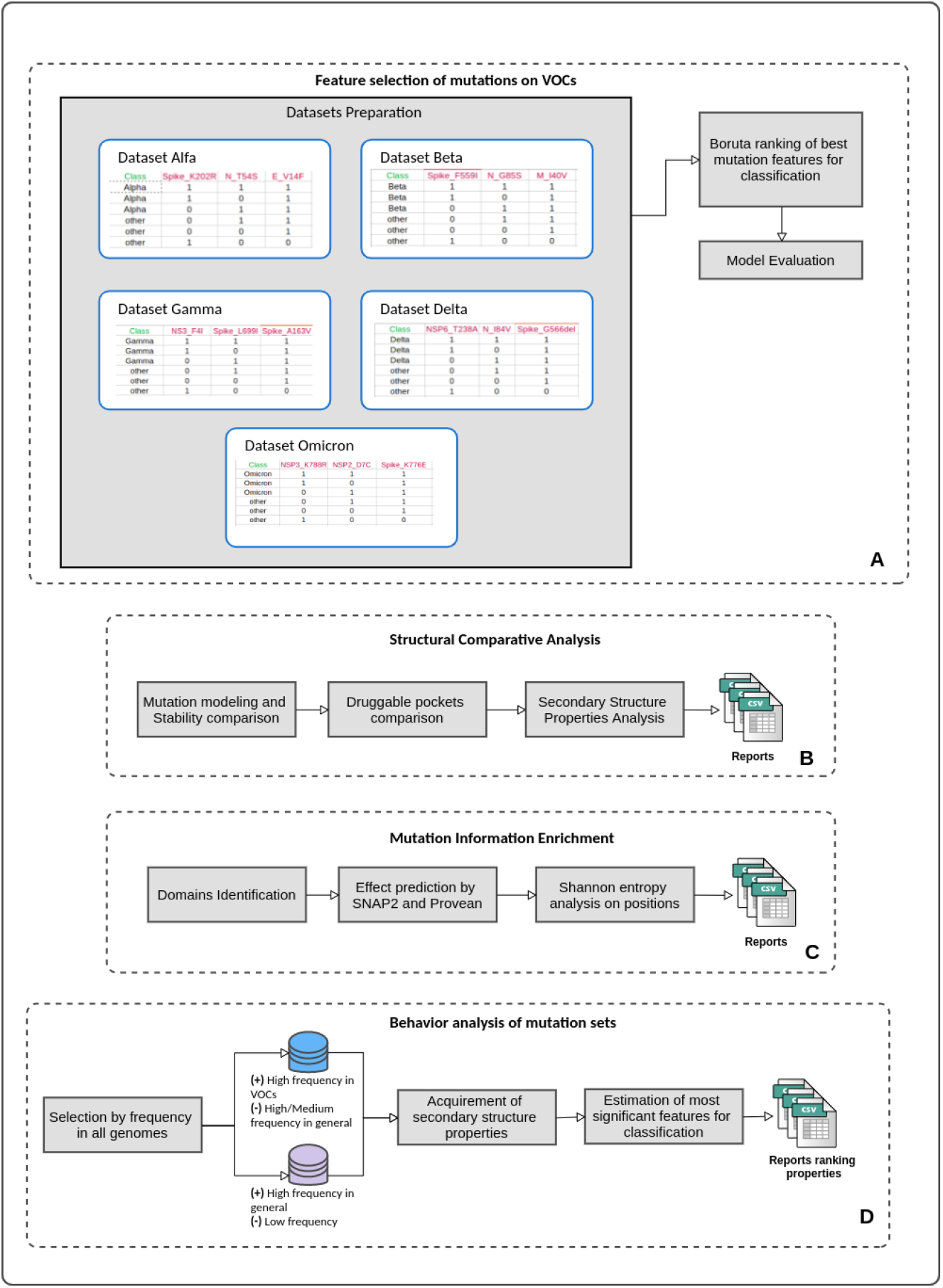
Diagram illustrating the steps of the following pipelines: (A) Feature selection of mutations in VOCs, (B) Structural Comparative analysis, (C) Mutation information enrichment, and (D) Behavior analysis of mutation sets. In the last pipeline (D), (+) represents the filter applied for the positive samples set while (-) represents the negative ones.

### 2.2 Structural comparative analysis pipeline

The structural comparative analysis is comprised of three main components (Figure 1 B): Structural modeling of the amino acid mutations with quality and stability analysis; comparative analysis of druggable pockets between reference and mutated protein; and secondary structure properties comparative analysis. The first component received as input the mutations in a specific chain and the reference protein structure, and used the Foldx tool ([3]) to build a structure model with the mutations whose positions were matched in the reference structure. Using this structure, the pipeline generated a Ramachandran plot ([12]) indicating the number of amino acid locations that were in a favorable region and checking the model quality. The last function of this component then evaluated the stability of reference and mutated structure by comparing the energy parameters provided by FoldX Stability function ([3]).

The second component executed the Fpocket software ([18]) for the reference and mutated protein, and evaluated the positions and druggable scores, returning the possible changes of binding sites promoted by the mutations.

The last component analyzed the reference primary sequence ([43]) and the one with mutations of interest by predicting the composition of secondary structure (helix, strand or coil), the solvent accessibility (bury, medium exposed or exposed) and the disorder of each position. All these components returned specific conformation changes promoted by the mutations and side chains altered by possible polarization provided by the charges of amino acid changes.

### 2.3 Mutation information enrichment

In order to enrich information about the significant mutations in the SARS-CoV-2 proteins, we developed a pipeline (Figure 1 C) to retrieve information from four data sources such as domains annotations from Prosite ([34]), functional impact prediction by SNAP2 ([11]) and a temporal series analysis of entropy ([32]).

The protein domains correspond to conserved regions responsible for a certain function, and depending on the mutation positions some characteristic domains of a protein may not be recognized promoting a loss of function. The first enrichment component annotated the SARS-CoV-2 proteins according to the Prosite tool ([34]) to identify the concentration of mutations in each domain region and the cases in which the domains were not identified.

The second component complements the domain information with the prediction of the functional impact of mutations of all proteins. We retrieved the reference primary sequences from Uniprot^1^ and applied all of them to SNAP2 ([11]) webserver retrieving the prediction scores and the classification of the impact (neutral or effect) for each position and possible amino acid change. The functional impact is also evaluated by PROVEAN ([5]) to support and complement the SNAP2 prediction.

The third component analyzed the shannon entropy ([32]) of specific protein positions to correlate the disorder degree with their functional and structural changes. Using the metadata provided by GISAID ([33]) for the samples, we built a time series dataset according to the collection date and the amino acid mutations, and calculated the frequency and absolute sum of aminoacid changes in the positions by month.

### 2.4 Analysis of feature patterns on mutation sets

This analysis (Figure 1 D) was divided in two groups, the first (purple dataset) concerned the exploration of possible features that differ the behavior of mutations occurring in more than 1% (positive samples) of the total number of genomes and mutations with frequency below 0.1% (negative samples). The second group (blue dataset) contains mutations found in VOCs with frequency above 90% (positive samples) and mutations with frequency between 0.05% and 1%, which correspond to frequent mutations but not fixed in the VOCs. The frequencies were calculated using the GISAID metadata release of January, 11 2022. The frequencies of changes for each position in Spike were organized considering all lineages and a separated table was also calculated for each VOC population of genomes. For each group, the datasets were formed using the mutations as samples (lines) and the features consist in extracting the number of changes caused by each individual mutation in the secondary structure, disorder and solvent accessibility provided by the reports returned in the structural analysis pipeline (Section 2.2), with addition of possible changes of functional group from the mutated position (ex. Positive Charge - Negative Charge). We use the Boruta method [16] to calculate the importance of each feature and rank them according to the performance on Random Forest classifier [28]. We perform a 10-fold cross validation to guarantee diversity and avoid overfitting the model. All the 10 datasets of each group had the same number of positive and negative samples. In the first group, the positive samples were the mutations with high frequency and the negative samples were the mutations with low frequency. In the second group, the positive samples were the mutations with high frequency in the VOCs and the negative samples, the mutations

## 3 Results & Discussion

The screening analysis of mutations present in the VOC aims the characterization of patterns by *in-silico* methods. All the 2021 and 2022 (up to January, 11) GISAID samples belonging to the five lineages that originated the VOCs were used for feature selection. Firstly, the filter considered only the mutations occurring in at least 80% of the samples of each lineage, then 126 shared mutations (intersection group, Supplementary table S1) for all VOCs were separated and excluded from the feature selection dataset, since these data columns would not contribute for the model. The feature selection analysis fixed four co-variables in the model (age, gender, lineage and country) identifying, in each re-sampling experiment the mutations ranked as 1 by Boruta were kept, at the end only the mutations that occurred in all re-sampling results remained for further analysis. The essential mutations discovered by the feature selection were grouped for each VOC.

The mutation enrichment component aggregated the information of functional impact predicted by SNAP and PROVEAN, the domain regions and the change of amino acid group, for the five sets of mutations and the intersection group (Supplementary Table S1-S6). According to this analysis, in each VOC group, mutations in at least 14 distinct proteins (Spike, NSP3, NSP10, NSP12, NSP13, NSP2, NSP4, NSP7, M, NSP14, NSP15, NSP16 and NSP9) contributed to differentiate one VOC from the others. For all groups, the major portion of mutations were derived from Spike, NSP12 and NSP3. These three proteins are essential for the infection in the host cell, replication and the host response. Spike interacts with the human protein ACE2 to enter the cell and start replication ([22]). NSP3 protein acts against the host inflammatory response by cleaving the poly-ubiquitin chains and reducing the type I interferon response ([2, 22]). NSP12 catalyzes the synthesis of viral RNA and, together with the non structural proteins NSP13-15, may help to antagonize the host innate immune response triggered by incoming viral RNAs or other signals at the beginning of infection ([22, 44]).

In all the five sets of mutations, the major part of the protein mutations (mainly in Spike and NSP3) was classified as neutral in both SNAP2 and predictors. Although certain proteins such as NSP10, NSP15 and NSP16 have a balanced number of mutations considered deleterious and having an effect. We observed that the proteins that concentrated high number of mutations classified as effect or deleterious, obtained in fact the smaller numbers of mutations, indicating a preference for proteins and positions in which the virus has less chances to be damaged.

We also assessed the secondary structural properties (Supplementary table S.7) of the mutations comprised in each of the five sets, and took into account only the positions whose properties have changed in the analysis in relation to the respective reference protein properties. Some patterns remained the same for all sets and proteins such as the major proportion of solvent accessibility reduction and more than 90% of disordered (less rigidity) amino acids position changing to ordered (less flexible). This change was previously described for the Spike protein ([45]) and the regions that occurs disordered to order transitions (DOTs) contribute to increase the binding energies for the spike-ACE2 complex formation. The report of secondary folding type changes showed that even mutations not occurring exactly in domain regions, the secondary folding was altered mainly in the Receptor Binding Domain (RBD) from Spike. The other proteins also presented these alterations in a balanced proportion of Strand, Coil and Helix pairwise combination. The P.1 set of mutations presented the major number of changes in the positions contained in RBD region, mainly from Helix to Strand. The changes of folding type also result in solvent accessibility alterations, the major number of position changes in relation to the reference proteins turned some regions to be buried such as the hydrophobic amino acids, tending to be found in the internal part of the 3D structure. The secondary structure properties analysis was previously reported for the SARS-CoV-2 genes considering a subset of amino acid mutations ([24]), our work expanded the analysis to all proteins and considered all the relevant mutations related to the VOCs. Interestingly, P.1 unique mutations as well as the shared mutations induced the highest numbers of changes in the solvent accessibility, increasing or decreasing this surface. While P.1 and shared mutations decreased turned 45 and 39 sites to bury or medium the other VOCs mutations obtained a maximum of 10, and Omicron changed zero sites. The unique mutations found for P.1 were also responsible for changing the major number of hot spots (17) that interacts with ACE2, only 2 of these affected sites increased exposure. In ascending order, the shared mutations and B.1.351 affected six hot spots, B.1.617.2 changed five sites, B.1.1.529 altered only one hot spot and B.1.1.7 promoted changes in zero sites. Since Spike has a great importance in the cell invasion mechanisms by binding to the human protein ACE2 and has been targeted for development of vaccines and antibodies evasion studies, we performed a detailed analysis focusing on the mutations found for this protein for the five VOCs.

### 3.1 Analysis of Spike significant mutations

For the Spike Protein, observing all mutations occurred in each of five variants of concern and their location in the Spike domains, the S1 NTD domain contains the most proportion of mutations, except in the omicron in which the highest proportion number occurred in the S1 CTD (Figure 2). The Gamma variant was the only VOC that presented a relevant number (around 10%) of mutations located in the S2 HR2 domain. The Gamma variant obtained the highest number of significant mutations (34), while the other variants reached a maximum of 12 (Figure 3). The significant mutations did not match any of the defining mutations of each variant. In order to investigate the energy parameters of the protein structure stability, we modeled the significant mutations in the reference Spike structure (PDB 6VXX) in two scenarios (Supplementary table S.8): using only the significant mutations, and using the variant defining plus significant mutations. The total energy of all the modeled structures presented negative variation, producing more stable structures. The highest negative variations occurred for the Omicron variant, while the energy variation with significant mutations only was -23.210 the addition of all its defining mutations helped to increase this number to -25.290. In P.1, the addition of the defining mutations produced an even higher stability increase, from -6.705 to -21.080.

**Fig. 2.**
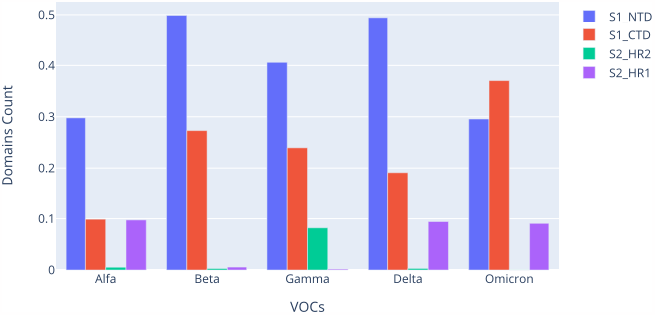
Distribution of the significant mutations along in the Spike domains.

**Fig. 3.**
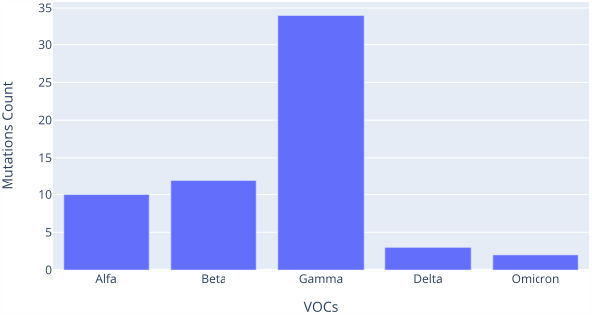
Distribution of the significant mutations along the variants.

The mutation W154C found significant to separate omicron from the others in NSP8 protein is responsible for changing the interface of this non structural protein. Together with the Tryptophan to Cysteine substitution W182C, produce an outlier number od variation of energy, suggesting that some backbone rearrangement is necessary in response to exchange of the large Tryptophan side chains for smaller Cysteines [21].

Among the shared mutations, the missense change T572I occurring in parallel with A570L, F855Y and N856I increased the mobility in RBD domain, showing that targeting the interdomain positions destabilizes the closed state, enhances the stability of the open form, and then may alters the dynamic preferences of the SARS-CoV-2 S trimer, according to conformational dynamics simulations [42]. Complementing this finding, the positions located in 570-572 and 856-862 may be involved in regulating functional motions near the interdomain regions and overlapped with positions used in protein design experiments in order to adjust the dynamic equilibrium and induce a population shift of the SARS-CoV-2 spike protein [40]. The shared mutation A653V, predicting according to our enrichment analysis as neutral by both SNAP and Provean, also stabilizes the Spike protein according to the stability predictors used by [23], besides this mutation cause loss of the hydrophobic core of the gear domain compactness [7]. Mutations located around the gear domain have also been proposed as drivers of ongoing viral adaptation [26]. The shared mutation H655Y together with N679K and P681H, present in the new variant of Concern Omicron, allow for the entry of viruses into cells and enhance the viral replication and infectivity [14]. According to a molecular dynamics simulations analysis, the shared V1176F mutation promotes the flexibility of the Spike protein by increasing motility and inducing compactness [13]. In addition, this mutation is also associated with higher patient mortality [9]. The shared mutation T716I, a change from Polar aa to a non-Polar one, was predicted as having an effect by SNAP and deleterious as provean, besides it is one of the defining mutations of Alfa variant, this mutations occurs in samples of the other variants with low frequency [30]. The shared mutation L5F that is located at the signal peptide is often found in sequences of all variants, although some work handle this mutation as a possible sequence processing error [15], other works report that this mutation increases hydrophobicity of the signal peptide, thus facilitate in viral secretion from cell and enhances epitope binding affinity [29]. Thus, the L5F is a disadvantageous for the SARS-CoV-2 virus since the mutated epitope could enhance CD8 T cell recognition and killing through the enhanced interaction with most of the HLA alleles [8].

According to docking analysis, the P.1 significant change N439K increases the affinity binding between Spike and hACE2 through the formation of a new salt bridge [13], the evaluation of clinical data and outcomes of viruses carrying this mutation also demonstrate the association with high fitness and virulence. Additionally, immunologic assays also show that mutations that produce equivalent viral viral fitness may induce immune evasion from both monoclonal and polyclonal antibody responses [38], affecting antibodies neutralization [39]. The Spike mutation S221L was predicted as a destabilizing mutation by recent works [23, 1], in fact the secondary structure prediction analysis showed that the 34 Spike mutations promote 71 changes of solvent accessibility properties: Bury to Exposed (4), Bury to Medium (16), Exposed to Bury (13), Exposed to Medium (12), Medium to Bury (20) and Medium to Exposed (6). The S221L mutation isolated promotes the change from exposed to bury in the 218 position [25]. The mutation E484A that have been also found in Omicron presents resistance to convalescent plasma along with E484K, E484G and E484D [19], and also evades the neutralizing antibody REGN10933 [41]. This behavior was also validated by assays [35, 20] showing that this mutation abolished binding of the monoclonal antibody 2B04, diminished binding by 2C03 but no effect was observed in relation to the monoclonal antibodies 2C02 or 2E06.

According to entropy analysis, the position 484 presented high variability every month (Figure 4), indicating that this position has been a hot spot for mutations. The Shannon entropy analysis also demonstrated that sixteen positions presented high variability in sixteen months or more (Figure 5). The position 614 rarely presents an alternative mutation to D (aspartic acid) to G (glycine) as well as another 1817 positions whose entropy value is less than 0.1.

**Fig. 4.**
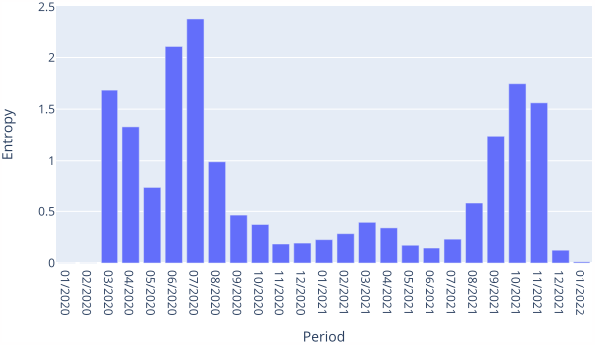
Shannon entropy values for Spike position 484 from 01/2020 to 01/2022.

**Fig. 5.**
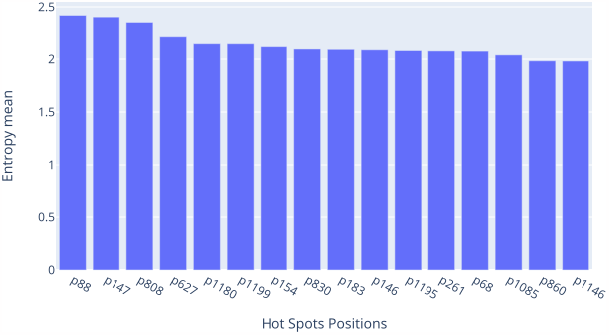
Shannon entropy values for Spike positions with entropy above 1.6 for sixteen months or more.

### 3.2 Analysis of feature patterns differentiating mutations with high from low frequency

Applying the filters of total frequency described in section 2.4, the first group of analysis were composed of 56 Spike mutations with high frequency (positive samples) and 1285 mutations with low frequency. The second group were composed of 37 Spike mutations with high frequency in VOCs (positive samples) and 223 mutations with low frequency. The ten rounds of the cross validation the indexes were randomly chosen to form distinct datasets.

According to the structural effects predicted by the proposed pipeline, we ran the stability and the secondary structure analysis functions and parsed the reports comparing the changes in each position induced by the mutation. The features names were composed by the pair of changed properties (ex. for solvent accessibility one of the possibilities may be Bury-Exposed). The value of these features are given by the sum of positions that obtained the same type of property change. Then, the features of the first group were: snap_prediction, B-Bury-M-Medium, E-Strand-H-Helix, .-ordered-*-disordered, E-Strand-C-Coil, Positive Charge-Polar, Polar-Non Polar, Non Polar-Polar, B-Bury-E-Exposed, M-Medium-E-Exposed, E-Exposed-B-Bury, M-Medium-B-Bury, H-Helix-E-Strand, C-Coil-H-Helix, Negative Charge-Non Polar, H-Helix-C-Coil, Polar-Positive Charge, E-Exposed-M-Medium, Negative Charge-Polar, *-disordered-.-ordered, C-Coil-E-Strand, Positive Charge-Non Polar, Non Polar-Negative Charge, Non Polar-Positive Charge, Negative Charge-Positive Charge. The features corresponding to the second group datasets were: snap_prediction, B-Bury-M-Medium, E-Strand-H-Helix, .-ordered-*-disordered, E-Strand-C-Coil, Positive Charge-Polar, Polar-Non Polar, Non Polar-Polar, B-Bury-E-Exposed, M-Medium-E-Exposed, E-Exposed-B-Bury, M-Medium-B-Bury, H-Helix-E-Strand, C-Coil-H-Helix, Negative Charge-Non Polar, H-Helix-C-Coil, E-Exposed-M-Medium, Polar-Positive Charge, Negative Charge-Polar, *-disordered-.-ordered, C-Coil-E-Strand, Non Polar-Negative Charge, Non Polar-Positive Charge, Negative Charge-Positive Charge.

The goal of this analysis is identifying a pattern of behavior according to these structural predicted properties that could identify important alterations that could alert an increase and preference for a given mutation in the virus replication. We took into consideration the features ranked in first and second places, for the first group the main features separating the two classes of samples corresponds mostly with changes in solvent accessibility from medium to bury, showing that this structural change may be considered crucial for the increase and prevalence of mutations. The other top ranked features were related to functional group change from polar to positive charge, contributing to change the electrostatic surface and energies stability of the protein structure. In the Second group the top ranked for most of the ten rounds was also the change of solvent accessibility previously observed (Medium to Bury), differently from the first group, the other top ranked feature was the change of secondary structure type from coil to strand.

[37] shows antigenic changes promoted by h3n2 and sars-cov-2, E484K reduced neutralization of many classes of antibodies, and is even worse combined with N501Y. Using the same features mentioned above, a clustering assay performed using K-means algorithm showed that in 100 simulations with mixed datasets, E484K always is grouped together with N501Y. The mutations L452R and P681H were also grouped together, L452 reduces antibodies neutralization and the S1/S2 furin cleavage site at position 681 promotes infection and cell-cell fusion [4]. PMC8114812 shows that H655Y and D796Y co occurs with mutations (L18F, L452R and N501Y) that promotes immune escape and/or higher infectivity and transmissibility, but these two mutations were not functionally described yet, and our clustering analysis also grouped them together. The mutations T716I and D1118H that were grouped in another cluster are both defining mutations of Alfa variant and together of the other mutations are assumed to be involved in the high infection rate [30]. The mutation S221L, found significant for Gamma variant, destabilizes the protein structure as well as 122 mutations, from these our clustering analysis grouped T95I, G1124V in the same cluster.

## Acknowledgments

We used GISAID sequence data to perform the analysis, the exactly sequences are detailed in the methods.

## Footnotes

1 http://www.uniprot.org

## References

[1] Reem Y Aljindan, Abeer M Al-Subaie, Ahoud I Al-Ohali, Balu Kamaraj, et al. Investigation of nonsynonymous mutations in the spike protein of sars-cov-2 and its interaction with the ace2 receptor by molecular docking and mm/gbsa approach. Computers in biology and medicine, 135:104654, 2021.

[2] Silvia Angeletti, Domenico Benvenuto, Martina Bianchi, Marta Giovanetti, Stefano Pascarella, and Massimo Ciccozzi. COVID-2019: The role of the nsp2 and nsp3 in its pathogenesis. J. Med. Virol., 92(6):584–588, June 2020.

[3] Oliver Buß, Jens Rudat, and Katrin Ochsenreither. Foldx as protein engineering tool: better than random based approaches? Computational and structural biotechnology journal, 16:25–33, 2018.

[4] Sébastien Calvignac-Spencer, Matthias Budt, Matthew Huska, Hugues Richard, Luca Leipold, Linus Grabenhenrich, Torsten Semmler, Max von Kleist, Stefan Kröger, Thorsten Wolff, et al. Rise and fall of sars-cov-2 lineage a. 27 in germany. Viruses, 13(8):1491, 2021.

[5] Yongwook Choi and Agnes P Chan. PROVEAN web server: a tool to predict the functional effect of amino acid substitutions and indels. Bioinformatics, 31(16):2745–2747, August 2015.

[6] Nuno R Faria, Thomas A Mellan, Charles Whittaker, Ingra M Claro, Darlan da S Candido, Swapnil Mishra, Myuki AE Crispim, Flavia CS Sales, Iwona Hawryluk, John T McCrone, et al. Genomics and epidemiology of the p. 1 sars-cov-2 lineage in manaus, brazil. Science, 372(6544):815–821, 2021.

[7] Jorge González-Puelma, Jacqueline Aldridge, Marco Montes de Oca, Mónica Pinto, Roberto Uribe-Paredes, José Fernández-Goycoolea, Diego Alvarez-Saravia, Hermy Álvarez, Gonzalo Encina, Thomas Weitzel, et al. Mutation in a sars-cov-2 haplotype from sub-antarctic chile reveals new insights into the spike’s dynamics. Viruses, 13(5):883, 2021.

[8] Elisa Guo and Hailong Guo. Cd8 t cell epitope generation toward the continually mutating sars-cov-2 spike protein in genetically diverse human population: Implications for disease control and prevention. PloS one, 15(12):e0239566, 2020.

[9] Georg Hahn, Chloe M Wu, Sanghun Lee, Julian Hecker, Sharon Lutz, Sebastien Haneuse, Dandi Qiao, Dawn DeMeo, Rudolph Tanzi, Manish Choudhary, et al. Two mutations in the sars-cov-2 spike protein and rna polymerase complex are associated with covid-19 mortality risk. 2021.

[10] Pengcheng Han, Chao Su, Yanfang Zhang, Chongzhi Bai, Anqi Zheng, Chengpeng Qiao, Qing Wang, Sheng Niu, Qian Chen, Yuqin Zhang, Weiwei Li, Hanyi Liao, Jing Li, Zengyuan Zhang, Heecheol Cho, Mengsu Yang, Xiaoyu Rong, Yu Hu, Niu Huang, Jinghua Yan, Qihui Wang, Xin Zhao, George Fu Gao, and Jianxun Qi. Molecular insights into receptor binding of recent emerging SARS-CoV-2 variants. Nat. Commun., 12(1):6103, October 2021.

[11] Maximilian Hecht, Yana Bromberg, and Burkhard Rost. Better prediction of functional effects for sequence variants. BMC Genomics, 16 Suppl 8:S1, June 2015.

[12] Rob WW Hooft, Chris Sander, and Gerrit Vriend. Objectively judging the quality of a protein structure from a ramachandran plot. Bioinformatics, 13(4):425–430, 1997.

[13] Szu-Wei Huang and Sheng-Fan Wang. Sars-cov-2 entry related viral and host genetic variations: Implications on covid-19 severity, immune escape, and infectivity. International Journal of Molecular Sciences, 22(6):3060, 2021.

[14] Yao Jiang, Qian Wu, Peipei Song, and Chongge You. The variation of sars-cov-2 and advanced research on current vaccines. Frontiers in Medicine, 8, 2021.

[15] Bette Korber, Will M Fischer, Sandrasegaram Gnanakaran, Hyejin Yoon, James Theiler, Werner Abfalterer, Nick Hengartner, Elena E Giorgi, Tanmoy Bhattacharya, Brian Foley, et al. Tracking changes in sars-cov-2 spike: evidence that d614g increases infectivity of the covid-19 virus. Cell, 182(4):812–827, 2020.

[16] Miron B Kursa, Aleksander Jankowski, and Witold R Rudnicki. Boruta–a system for feature selection. Fundamenta Informaticae, 101(4):271–285, 2010.

[17] Miron B Kursa, Witold R Rudnicki, et al. Feature selection with the boruta package. J Stat Softw, 36(11):1–13, 2010.

[18] Vincent Le Guilloux, Peter Schmidtke, and Pierre Tuffery. Fpocket: an open source platform for ligand pocket detection. BMC bioinformatics, 10(1):1–11, 2009.

[19] Xue Li, Liying Zhang, Si Chen, Weilong Ji, Chang Li, and Linzhu Ren. Recent progress on the mutations of sars-cov-2 spike protein and suggestions for prevention and controlling of the pandemic. Infection, Genetics and Evolution, 93:104971, 2021.

[20] Zhuoming Liu, Laura A VanBlargan, Louis-Marie Bloyet, Paul W Rothlauf, Rita E Chen, Spencer Stumpf, Haiyan Zhao, John M Errico, Elitza S Theel, Mariel J Liebeskind, et al. Identification of sars-cov-2 spike mutations that attenuate monoclonal and serum antibody neutralization. Cell host & microbe, 29(3):477–488, 2021.

[21] Joseph H Lubin, Christine Zardecki, Elliott M Dolan, Changpeng Lu, Zhuofan Shen, Shuchismita Dutta, John D Westbrook, Brian P Hudson, David S Goodsell, Jonathan K Williams, et al. Evolution of the sars-cov-2 proteome in three dimensions (3d) during the first 6 months of the covid-19 pandemic. Proteins: Structure, Function, and Bioinformatics, 2021.

[22] Giuseppina Mariano, Rebecca J Farthing, Shamar L M Lale-Farjat, and Julien R C Bergeron. Structural characterization of SARS-CoV-2: Where we are, and where we need to be. Front Mol Biosci, 7:605236, December 2020.

[23] Taj Mohammad, Arunabh Choudhury, Insan Habib, Purva Asrani, Yash Mathur, Mohd Umair, Farah Anjum, Alaa Shafie, Dharmendra Kumar Yadav, Md Hassan, et al. Genomic variations in the structural proteins of sars-cov-2 and their deleterious impact on pathogenesis: A comparative genomics approach. Frontiers in cellular and infection microbiology, p. 951, 2021.

[24] Thanh Thi Nguyen, Pubudu N Pathirana, Thin Nguyen, Quoc Viet Hung Nguyen, Asim Bhatti, Dinh C Nguyen, Dung Tien Nguyen, Ngoc Duy Nguyen, Douglas Creighton, and Mohamed Abdelrazek. Genomic mutations and changes in protein secondary structure and solvent accessibility of SARS-CoV-2 (COVID-19 virus). Sci. Rep., 11(1):3487, February 2021.

[25] Thanh Thi Nguyen, Pubudu N Pathirana, Thin Nguyen, Quoc Viet Hung Nguyen, Asim Bhatti, Dinh C Nguyen, Dung Tien Nguyen, Ngoc Duy Nguyen, Douglas Creighton, and Mohamed Abdelrazek. Genomic mutations and changes in protein secondary structure and solvent accessibility of sars-cov-2 (covid-19 virus). Scientific Reports, 11(1):1–16, 2021.

[26] Thomas P Peacock, Rebekah Penrice-Randal, Julian A Hiscox, and Wendy S Barclay. Sars-cov-2 one year on: evidence for ongoing viral adaptation. The Journal of general virology, 102 (4), 2021.

[27] Arun K Pujari. Data mining techniques. Universities press, 2001.

[28] Yanjun Qi. Random forest for bioinformatics. In Ensemble machine learning, pp. 307–323. Springer, 2012.

[29] Md Marufur Rahman, Shirmin Bintay Kader, and SM Shahriar Rizvi. Molecular characterization of sars-cov-2 from bangladesh: Implications in genetic diversity, possible origin of the virus, and functional significance of the mutations. Heliyon, 7(8):e07866, 2021.

[30] Renuka Raman, Krishna J Patel, and Kishu Ranjan. Covid-19: Unmasking emerging sars-cov-2 variants, vaccines and therapeutic strategies. Biomolecules, 11(7):993, 2021.

[31] Sindhu Ramesh, Manoj Govindarajulu, Rachel S Parise, Logan Neel, Tharanath Shankar, Shriya Patel, Payton Lowery, Forrest Smith, Muralikrishnan Dhanasekaran, and Timothy Moore. Emerging SARS-CoV-2 variants: A review of its mutations, its implications and vaccine efficacy. Vaccines (Basel), 9(10), October 2021.

[32] C E Shannon. A mathematical theory of communication. The Bell System Technical Journal, 27(3):379–423, July 1948.

[33] Yuelong Shu and John McCauley. Gisaid: Global initiative on sharing all influenza data–from vision to reality. Eurosurveillance, 22(13):30494, 2017.

[34] Christian J A Sigrist, Lorenzo Cerutti, Edouard de Castro, Petra S Langendijk-Genevaux, Virginie Bulliard, Amos Bairoch, and Nicolas Hulo. PROSITE, a protein domain database for functional characterization and annotation. Nucleic Acids Res., 38(Database issue):D161–6, January 2010.

[35] Wen Su, Sin Fun Sia, Aaron J Schmitz, Traci L Bricker, Tyler N Starr, Allison J Greaney, Jackson S Turner, Bassem M Mohammed, Zhuoming Liu, Ka Tim Choy, et al. Neutralizing monoclonal antibodies that target the spike receptor binding domain confer fc receptor-independent protection against sars-cov-2 infection in syrian hamsters. Mbio, 12(5):e02395–21, 2021.

[36] Kaiming Tao, Philip L Tzou, Janin Nouhin, Ravindra K Gupta, Tulio de Oliveira, Sergei L Kosakovsky Pond, Daniela Fera, and Robert W Shafer. The biological and clinical significance of emerging SARS-CoV-2 variants. Nat. Rev. Genet., September 2021.

[37] Houriiyah Tegally, Eduan Wilkinson, Marta Giovanetti, Arash Iranzadeh, Vagner Fonseca, Jennifer Giandhari, Deelan Doolabh, Sureshnee Pillay, Emmanuel James San, Nokukhanya Msomi, et al. Detection of a sars-cov-2 variant of concern in south africa. Nature, 592(7854):438–443, 2021.

[38] Emma C Thomson, Laura E Rosen, James G Shepherd, Roberto Spreafico, Ana da Silva Filipe, Jason A Wojcechowskyj, Chris Davis, Luca Piccoli, David J Pascall, Josh Dillen, et al. Circulating sars-cov-2 spike n439k variants maintain fitness while evading antibody-mediated immunity. Cell, 184(5):1171–1187, 2021.

[39] Norma A Valdez-Cruz, Enrique García-Hernández, Clara Espitia, Laura Cobos-Marín, Claudia Altamirano, Carlos G Bando-Campos, Luis F Cofas-Vargas, Enrique W Coronado-Aceves, Ricardo A González-Hernández, Pablo Hernández-Peralta, et al. Integrative overview of antibodies against sars-cov-2 and their possible applications in covid-19 prophylaxis and treatment. Microbial cell factories, 20(1):1–32, 2021.

[40] Gennady M Verkhivker. Molecular simulations and network modeling reveal an allosteric signaling in the sars-cov-2 spike proteins. Journal of proteome research, 19(11):4587–4608, 2020.

[41] Gennady M Verkhivker, Steve Agajanian, Deniz Yazar Oztas, and Grace Gupta. Comparative perturbation-based modeling of the sars-cov-2 spike protein binding with host receptor and neutralizing antibodies: Structurally adaptable allosteric communication hotspots define spike sites targeted by global circulating mutations. Biochemistry, 60(19):1459–1484, 2021.

[42] Gennady M Verkhivker and Luisa Di Paola. Dynamic network modeling of allosteric interactions and communication pathways in the sars-cov-2 spike trimer mutants: Differential modulation of conformational landscapes and signal transmission via cascades of regulatory switches. The Journal of Physical Chemistry B, 125(3):850–873, 2021.

[43] Sheng Wang, Wei Li, Shiwang Liu, and Jinbo Xu. RaptorX-Property: a web server for protein structure property prediction. Nucleic Acids Res., 44(W1):W430–5, July 2016.

[44] Wenjing Wang, Zhuo Zhou, Xia Xiao, Zhongqin Tian, Xiaojing Dong, Conghui Wang, Li Li, Lili Ren, Xiaobo Lei, Zichun Xiang, and Jianwei Wang. SARS-CoV-2 nsp12 attenuates type I interferon production by inhibiting IRF3 nuclear translocation. Cell. Mol. Immunol., 18(4):945–953, April 2021.

[45] Dhanusha Yesudhas, Ambuj Srivastava, Masakazu Sekijima, and M Michael Gromiha. Tackling covid-19 using disordered-to-order transition of residues in the spike protein upon angiotensin-converting enzyme 2 binding. Proteins, 89(9):1158–1166, September 2021.

